# Predicting Clinical Outcomes in Infants With Cystic Fibrosis From Stool Microbiota using Random Forest Algorithms

**DOI:** 10.1101/2022.08.06.503028

**Authors:** Rebecca A. Valls, Thomas H. Hampton, Courtney E. Price, Kaitlyn E. Barrack, George A. O’Toole, Modupe O. Coker, Juliette C. Madan

**Author notes:** **Correspondence:** Modupe Coker, **Juliette Madan**.

## Abstract

The gut-lung axis describes the interaction between microbiota in the gut and health status of the airway, whereby there is a bidirectional relationship facilitated by systemic transport of microbially-derived metabolites and immune factors. Cystic fibrosis (CF) is a genetic disease that is associated with dysbiosis of the gut microbiota. Recent literature has shown that the microbial dysbiosis in the CF gut can alter the hosts’ inflammatory status and that there are distinct microbial compositions in children with CF who have low versus high intestinal inflammation. These distinct microbial profiles can be used as signatures in children with CF (cwCF) to predict health outcomes. Here, we use supervised machine learning to train a random forest model on the distinct microbial composition of cwCF to predict: age (as a validation of the method), frequency of upper respiratory infection (URIfreq), and neutrophil to lymphocyte ratio (NLR), a clinical marker for systemic inflammation that negatively correlates with lung function. We find that the out of bag error, a measure of model accuracy, is lower when predicting age for cwCF compared to children without CF, consistent with previous data. We are able to predict high URIfreq with only 16% error and high NLR with 27% error. This machine learning pipeline may allow physicians and microbiome researchers to use the stool microbiota of cwCF as a tool for identifying individuals with the more negative airway clinical outcomes from this population, and potentially allow for early intervention.

**Importance:** Children with CF (cwCF) often experience chronic respiratory infections, leading to progressive, irreversible lung function decline and significant morbidity and premature mortality. Modulator therapy has revolutionized the treatment of eligible adults with CF. cwCF as young as age 6 are now eligible to receive modulator therapy, although by early childhood many children have already experienced pulmonary exacerbations. Here we show that for cwCF, stool microbiota composition is associated with higher upper respiratory infection frequency and increased systemic inflammation. Our findings may aid in developing diagnostic tools that can allow physicians further understanding of which intestinal microbiota profiles are associated with health outcomes and to identify targets for preventative treatment for cwCF.

## Introduction

Cystic fibrosis (CF) is a genetic disease associated with chronic lung infections that lead to overall decline in lung health^1,2^. These chronic lung infections begin in early childhood and are often preceded by pulmonary exacerbation event(s). Pulmonary exacerbations in CF are nonstandardized events most commonly described as acute inflammatory responses to infection that lead to a sharp decrease in lung function measured by FEV1 (forced expiratory volume in 1 second)^1^. Although early research focused on the microbial communities in the lungs of children with CF (cwCF), recent research has shown connections between CF and changes in intestinal microbial communities, including in children ^3–8^. Such observations are not surprising, because CF alters the intestinal environment the microbial communities inhabit, affecting the microbes’ ability to establish themselves and flourish ^1,2,6,9^. Intestinal microbial communities are further modulated by the treatments that cwCF regularly undergo, such as: high calorie diet, frequent use of antimicrobial agents, pancreatic insufficiency, requirement for enzyme replacement, and most recently, CFTR modulators^3,5,7^. CFTR modulators like Ivacaftor not only improve lung function, but have recently been shown to return the CF intestinal environment to a more nonCF-like state by increasing pH and reducing viscosity of secreted mucin ^5^. Additionally, changes in microbial communities are associated with differences in local and systemic markers of inflammation^104^.

Previously reported changes in the gut microbiota in cwCF include reduced *Bifidobacterium* and *Bacteroides*, with an increase in *Escherichia-Shigella* and *Clostridium* – although these shifts can depend on the cohort being examined ^1,4,7^. Alpha diversity in cwCF increases as children age but is consistently lower than infants without CF ^1,2^. The changes in microbial communities outlined here are also likely associated with alterations in lung health outcomes ^11^. The communication between the microbial communities in the gut and the lungs is referred to as the “gut-lung axis”, whereby microbes in the gut stimulate both local and distal immune responses through their secretion of metabolites and other microbial factors ^1^. This interaction can be positive, for example by downregulating inflammation through the secretion of short chain fatty acids by these microbes, including butyrate, which can travel systemically and engage with free fatty acid receptors in the lungs ^12^. Additionally, *Bacteroides*-secreted glycolipids can activate colonic dendritic cells that induce local and systemic cytokine release, which in turn enhance resistance to pulmonary viral infection ^13^. Commensal derived metabolites, such as riboflavin, can also imprint the abundance of mucosal-associated invariant T cells ^14^. The interaction between gut microbiota and the host can also be negative, whereby bacteria can secrete virulence factors^1^. One example of a secreted virulence factor is the secretion of a proteolytic enteroxin, *B. fragilis* toxin (BFT). This toxin, secreted by *Bacteroides fragilis*, can cause diarrhea and intestinal inflammation through its activation of NF-kappa B pathways by triggering cleavage of E cadherin in host colonic epithelial cells ^15^.

These interactions between gut microbiota and the host have been shown to be especially important in the first few years of a child’s life, when communication between immune cells and intestinal microbiota trains the immune system and shapes how cwCF respond to future infections ^6,11,14,16,17,18^. Individuals with chronic lung diseases, such as asthma, emphysema, and CF have benefited from gut microbiome manipulation: it has previously been shown in both mouse models and clinical trials that manipulating the gut microbiome, whether by probiotics, prebiotics, or fecal microbial transplants, lung disease progression was attenuated ^19,20,21,22^.

Machine learning is increasingly being used with clinical data to assist disease diagnosis^23^. In many cases, knowledge gained from machine learning can both assist physicians in making a diagnosis and help with design of patient-specific treatments. For example, such predictive approaches have been used to design treatment based on early signs of diabetes and Covid-19 ^23–25^. Here, we applied a supervised machine learning pipeline to predict lung health outcomes in cwCF from stool microbiota composition, and we chose to use a random forest model because of its ease of interpretability.

## Results

### Random forest models can predict age of cwCF from stool microbiome composition

To test whether clinical outcomes can be predicted from stool microbiome composition data, 282 fecal samples were collected from 39 cwCF (**Table 1**) and 16S rRNA gene amplicon libraries were sequenced from these samples. To balance the samples per age group and to facilitate their training via a random forest model, samples were randomly subsetted into n=21 per age group. Relative abundances of stool microbiota were set as features for each sample. Each sample was further labeled with the age of cwCF at time of collection. The relative abundances of fecal microbiota were used to train a random forest model to predict the age category of cwCF (**Figure 1A,B**). The predictive power, or accuracy, of the model was determined from the out of bag score (OOB), a measure of how well a model predicted the age of a stool microbiome sample where age is not specified. When training a model with classified samples, our random forest model randomly selects 2/3 of all samples to build the decision trees that make the predictions, by default. The remaining samples (1/3) are not included in the training process. These “out-of-bag” (OOB) samples are used to validate the predictive power of the model. The OOB score reflects the number of correctly predicted sample classifications from the out of bag samples. In this case, the random forest predictive model correctly predicted age category of cwCF with 45% error (**Figure 1C)**, an error value higher than previously reported for infants without CF ^8^. This decrease in prediction accuracy is not surprising, as several studies have shown that there is a marked dysbiosis in the gut of cwCF^1,2,4,6,9,21^. Additionally, Hayden et al. showed that cwCF have a delayed maturation of gut microbiota ^8^. Thus, our data are consistent with previous findings in children, and indicate that the gut dysbiosis in CF may interfere with age-related community development in such a way that it makes predictions more difficult to make accurately.

**Table 1.** Patient data.

|  |  | N of Classification Groups<sup>a</sup><br>(patients/samples) |  |  |  |  |  |
| --- | --- | --- | --- | --- | --- | --- | --- |
|  |  | <1 | 1 | 2 | 3 | 4 | Sum |
| Age | All | 27/44 | 32/89 | 25/71 | 23/57 | 19/21 | 39/282 |
|  | Subset<sup>b</sup> | 19/21 | 15/21 | 14/21 | 14/21 | 15/21 | 36/105 |

|  |  | N of Classification Groups<br>(patients/samples) |  |  |  |
| --- | --- | --- | --- | --- | --- |
|  |  | Low | Medium | High | Sum |
| URIfreq |  | 8/73 | 8/68 | 12/105 | 28/246 |
| NLR |  | 6/11 | 8/11 | 7/11 | 14/33 |

<sup>a</sup>Patient and sample n per classification group.
<sup>b</sup>Subset of age indicates number of samples used to train the random forest model.

**Figure 1.**
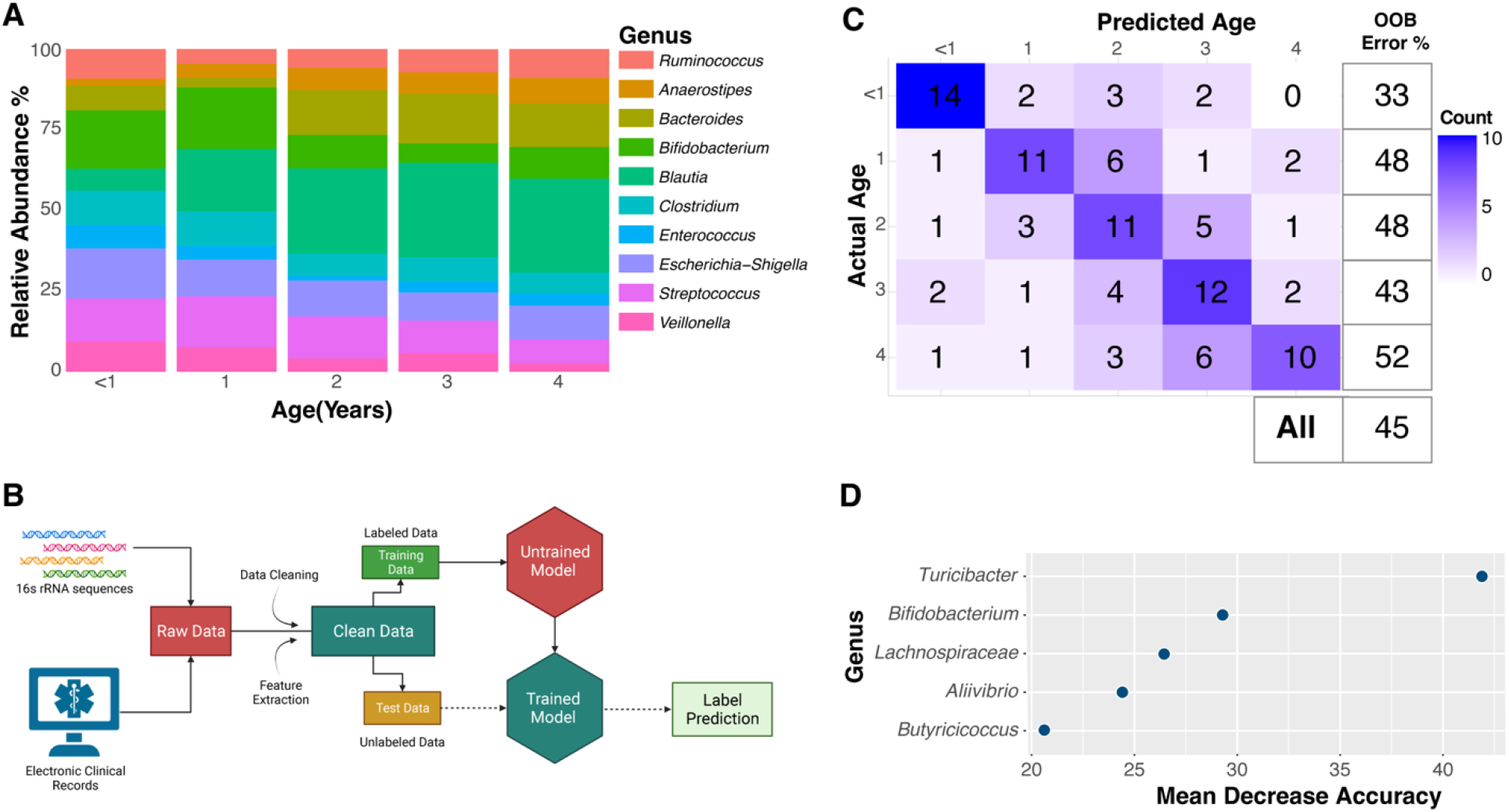
Random Forest Model for Predicting Age from Stool Microbiota. (A) Normalized relative abundance of the top 10 genera (~64% of total counts) in the stool samples collected from cwCF over the first four years of life. (B) Illustration of the pipeline for supervised machine learning through classification. Samples are labeled with their age and the normalized relative abundance of stool microbiota to train a random forest model to generate a predictive model for age. (C) Confusion matrix heatmap for the age prediction model. Overall OOB estimate of error rate for predicting age from stool microbiota is 45%. OOB is also shown for each indicated age. (D) Impact Factor summary plot for the top 5 stool microbiota important for classifying samples in the age model. *Turicibacter*, *Bifidobacterium*, and *Lachnoclostridium* are the top genera important for accurately predicting *age* of infant at which stool sample was collected.

The random forest model resulted in outputs for two measurements of importance. The Mean Decrease Accuracy, labeled as Impact Factor, lists the top features; in this case genera of stool bacteria, important for age-model output accuracy. For predicting age, these genera are *Turicibacter, Bifidobacterium*, and *Lachnoclostridium*. That is, if any of these genera were removed when generating the predictive model, the accuracy of the prediction would suffer. Although multiple iterations of the model resulted in a change in the order of the genera listed in **Figure 1D**, *Turicibacter* was consistently the top genus of bacteria driving the model output. A full list of the genera important for age-model output are shown in **Supplemental Figure 1A**. The second measure of importance to the model, Mean Decrease Gini, is a measure of node purity. The greater a feature’s Mean Decrease Gini, the better that genus is able to split classification groups into pure class nodes. For predicting age, *Bifidobacterium, Clostridium*, and *Ruminoccous* are the genera with highest node purity (**Supplemental Figure 1B**).

We next determined whether the genus *Turicibacter*, important for predicting age in cwCF, might inform other clinical outcomes. Specifically, we wondered whether the total number of URIs experienced by a patient would correlate with the relative abundance of *Turicibacter, Bifidobacterium*, and *Lachnoclostridium* in the stool across the first two years of life (**Figure 2A-C**). We focused on the first two years of life because it is during these early years that the gut microbiome is establishing; the gut microbiota is considered more stable after ages 3-5 ^1^, although some studies suggest earlier stabilization, as early as 1-3 years^3^. We found a modest, but statistically significant, positive correlation between total URIs and relative abundance of *Turicibacter* in stool (**Figure 2A**). Interestingly, 13 patients had no *Turicibacter* in the first two years of life (**Figure 2A**). These 13 patients were further assessed for correlations between total URI and relative abundance of *Bifidobacterium* or *Lachnoclostridium*, the two other genera contributing to model accuracy, to determine if another top age predictor could be driving total URI in cwCF lacking *Turicibacter*; no such correlation was identified. No statistically significant correlation was identified between total URI and relative abundance of *Bifidobacterium* or *Lachnoclostridium* (**Figure 2B,C**).

**Figure 2.**
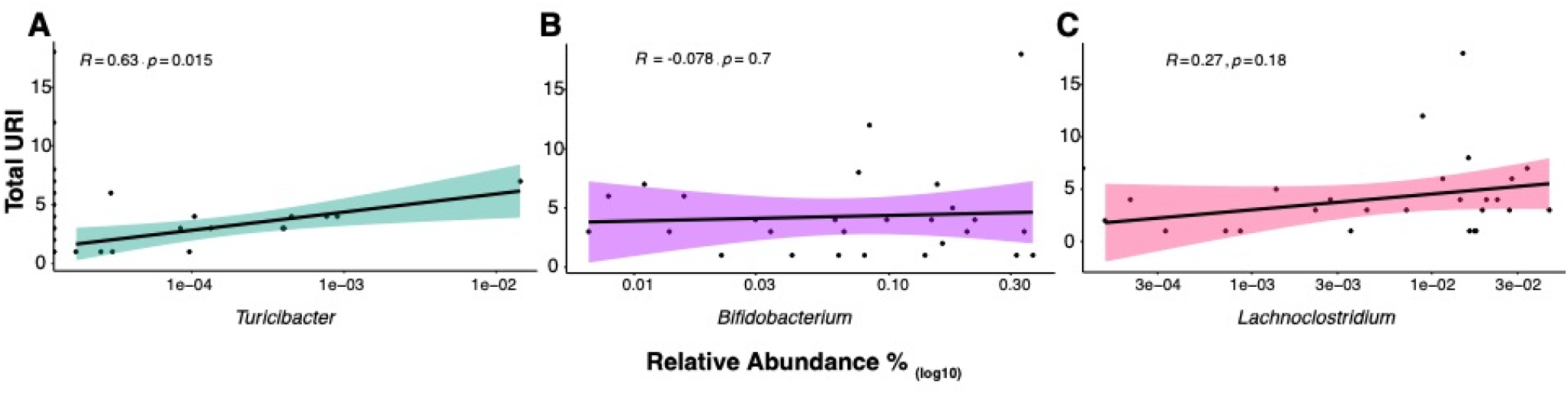
Correlating microbes and total URI. Correlation between total URI and log_10_ average relative abundance of (A) *Turicibacter*, (B) *Bifidobacterium*, and (C) *Lachnoclostridium* in the first two years of life. Each dot represents a single patient. *Turicibacter* shows significant positive correlation with Total URI when *Turicibacter* is present, but *Bifidoabcterium* and *Lachnoclostridium* do not. Spearman correlation performed with R and the p value is indicated.

### High URI frequency can be predicted for cwCF from stool microbiome composition

The modest correlation between *Turicibacter* and URI frequency suggested to us that directly applying random forests to the problem of predicting URI might better reveal specific taxa. Therefore, the frequency of upper respiratory infections (URIfreq) was calculated for each infant, that is, their total URI divided by age in years. We then stratified the infants into one of three URIfreq groups: Low (<1 URI per year of life), Medium (1 URI per year of life), and High (>1 URI per year of life). This stratification produced roughly equal numbers of cwCF in each group (**Table 1**).

Although the URIfreq predictive model predicted URIfreq OOB with 40% error overall, the model predicting high URIfreq only had 16% error (**Figure 3A**). The top genera of stool bacteria important for URIfreq-model output are *Faecalibacterium*, *Butyricioccus* and *Bacteroides* (**Figure 3B**). A full list of the genera important for URIfreq-model output are shown in **Supplemental Figure 2A, B**. *Faecalibacterium* and *Butyricioccus* secrete butyrate, a short chain fatty acid previously shown to reduce inflammation in the intestinal mucosa and be overall protective of gut physiology ^1,12,19^. Plotting relative abundance of the 10 most abundant stool microbiota across URIfreq groups revealed that *Bacteroides* doubled in the High URIfreq group compared to the Low and Medium URIfreq groups (**Figure 3C**), a point we address further in the Discussion. In order to determine whether a broader classification of microbiota could help inform function of microbes driving the predictive power of the URIfreq-model, we assessed the same model at a family level and found that the error did not change. The top family important for URIfreq prediction was *Butyricicoccaceae* (**Supplemental Figure 2C**).

**Figure 3.**
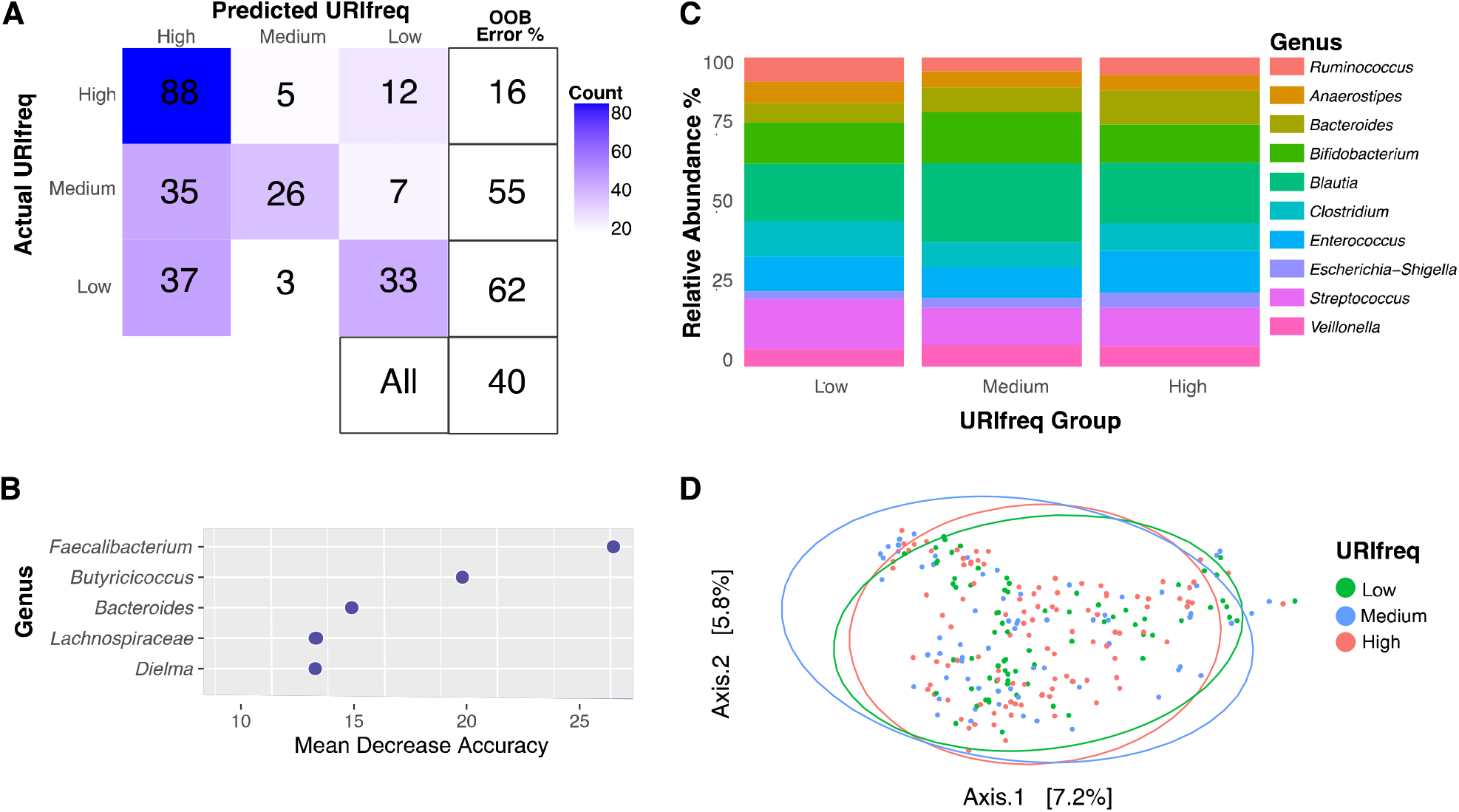
Association of Stool Microbiota with URI. (A) Confusion matrix heatmap for the URIfreq prediction model. The overall OOB estimate of error rate for predicting age from stool microbiota is 40%. Error rate for predicting High URIfreq is 16%. (B) The Impact Factor summary plot for the top 5 stool microbiota important for classifying samples in the age model. *Faecalibacterium*, *Butyricoccus*, and *Bacteroides* are the top genera important for accurately predicting URIfreq classification of cwCF. (C) Normalized relative abundance of top 10 genera (~64% of total counts) in the stool samples collected from infants with CF for each URIfreq classification group. (D) PCoA depicting beta diversity distances between URIfreq groups. Differences in community structure assessed by Bray-Curtis dissimilarity was revealed by PERMANOVA (adonis test) as a function of URIfreq group (P=0.001). However, factors other than URIfreq seem to drive most of the differences in composition because URIfreq explains little of the variability in composition (R2 = 0.016).

We next examined the microbial communities of the children classified as Low, Medium or High in upper respiratory infection frequency. The Chao1-determined alpha diversity, a measure of microbial richness, showed a significant increase in the high URIfreq group (**Supplemental Figure 3A**), but there was no significant increase in Shannon alpha diversity (**Supplemental Figure 3B**). This finding indicates that sample richness was different between the URIfreq groups, but not evenness. Microbial communities in the stool from children with different rates of upper respiratory infection frequency were substantially similar in composition. Statistically, significant differences in community structure assessed by Bray-Curtis dissimilarity was revealed by PERMANOVA (adonis test) as a function of URIfreq group (P=0.001). However, differences were very small and account for less than 2% of the variability (R2=0.016) (**Figure 3D**). When the interaction of age is accounted for, composition of community structure remains similar. Although statistically significant, (P=0.009), accounting for URIfreq group and age explain little of the variability in composition (R2=0.011) as determine by PERMANOVA (**Supplemental Figure 3C)**. Collectively, these results indicate that the overall microbial composition of each URIfreq group are similar.

### High NLR Classification can be predicted for cwCF from stool microbiome composition

We were able to obtain hematology data for a subset of cwCF in our cohort. Included in these hematology data were values for absolute neutrophil and lymphocyte levels. The ratio of absolute neutrophil count to absolute lymphocytes (NLR) provides a quantitative biomarker of systemic inflammation, where higher NLR has been shown to correlate with lower lung function (ppFEV_1_) ^27^. We therefore created random forest models to identify features in the CF infant stool microbiome composition that predict NLR.

As before, we began by stratifying our cwCF cohort into three groups: Low (0.2-0.46), Medium (0.47-0.85), or High (0.86-4) NLR, yielding an approximately equal number of individuals per group (**Table 1**). Although the overall OOB error of the NLR predictive model was 64%, the model performed well at predicting high NLR with 27% OOB error (**Figure 4A)**. The top genera of stool bacteria important for NLR prediction were *Lactococcus, Kluyvera*, and *Klebsiella* (**Figure 4B, Supplemental Figure 4A, B**). We saw no significant difference in the measure of alpha diversity (**Supplemental Figure 5)**. Statistically different community structure assessed by Bray-Curtis dissimilarity was revealed by PERMANOVA (adonis test) as a function of NLR groups. However, again this difference was very small and accounts for less than 10% of the variability (R2 = 0.085) in beta diversity among NLR groups in infants with CF (**Figure 4D**).

**Figure 4.**
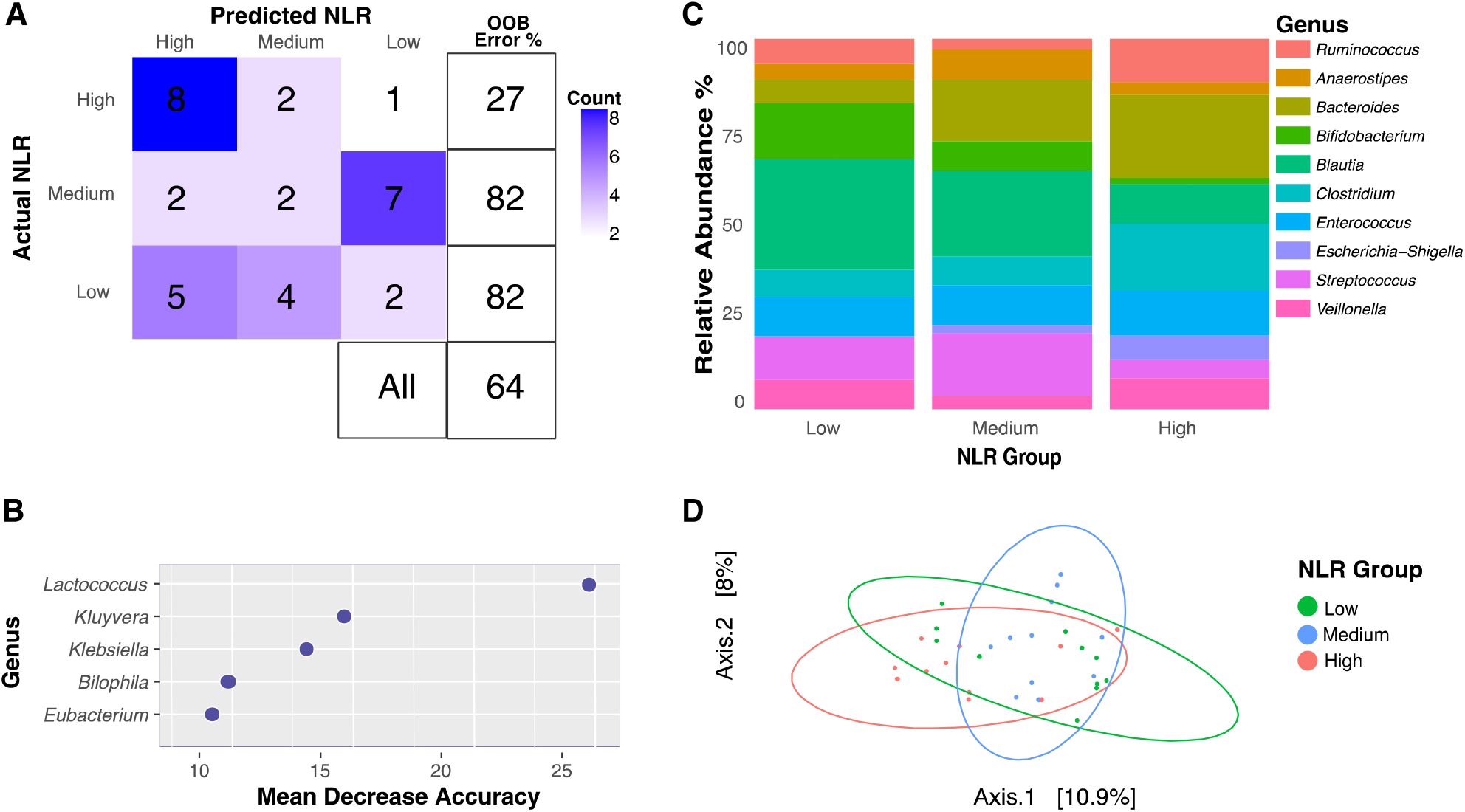
Association of Stool Microbiota with NLR. (A) Confusion matrix heatmap for the NLR prediction model. The overall OOB estimate of error rate for predicting NLR from stool microbiota is 64% but error rate for predicting High NLR is 27%. (B) Impact Factor summary plot for the top 5 genera important for classifying samples in the NLR model. *Lactococcus* is the top genus important for accurately predicting NLR classification group in pwCF. (C) Normalized relative abundances of top 10 genera (~64% of total counts) in the stool samples collected from infants with CF for each NLR classification group. (D) PCoA depicting beta diversity distances between NLR groups. Statistically different community structure assessed by Bray-Curtis dissimilarity was revealed by PERMANOVA (adonis test) as a function of NLR groups. Composition of NLR groups is systematically different (P==0.008). However, factors other than NLR seem to drive most of the differences in composition because NLR explains little of the variability in composition (R2 = 0.085) as determined by PERMANOVA.

## Discussion

This study used longitudinal stool and clinical data collections to train a model that can predict respiratory outcomes for children with CF from stool samples. Overall, the relative abundance of stool microbiota at the genus level in children with CF weakly predicts age, consistent with studies showing intestinal microbiome dysbiosis and altered age of maturation in this population^1,2,4,6,8,9,11,20,21,26,28^. Specifically, weak predictive power in cwCF compared to the general population suggests that the CF infant microbiome may mature at different rates than in infants without CF. Nonetheless, the genus of bacteria most important for predicting age in cwCF was *Turicibacter*. In a previous report from Hayden and colleagues^8^, *Blautia* was identified as a genus important for model accuracy for age prediction. In our analysis, *Blautia* also was identified as an important genus for predicting age prior to sub-sampling to balance the numbers of cwCF in each of the age groups. Upon subsampling, so that an even 21 samples per age group were used to train the model, *Turicibacter* was the top genus important for predicting age in infants with CF. However, we did not identify *Blautia* as a top genus for predicting age after subsampling. The Hayden Cohort^8^ consisted of cwCF ages 3 to 12 months. From this, we gather that *Blautia* may be important for predicting age in children in the first year of life but less so post 12 months of age. The genera *Blautia* and *Turicibacter* have also been identified as top contributors for predicting age from stool microbiota in children with asthma in the first year of life^18^.

In addition to its importance for predicting age, in our analysis *Turicibacter* was also found to have a positive correlation with total URI. *Turicibacter* has previously been shown to correlate with a decrease in butyrate in the CF gut ^21^, is considered a top contributor of acetate ^18^, and has been shown to positively correlate with inflammatory markers in asthmatic individuals ^29^. Thus, for the subset of individuals with *Turicibacter* (in only a subset of cwCF was this genus detected), this microbe could serve as a biomarker to calibrate “microbiota age” and perhaps help predict the frequency of negative respiratory outcomes.

Of clinical relevance, applying a random forest model to the stool microbiome of infants with CF could also predict cwCF in the “high” URIfreq group with relatively low error. The model’s predictive power for the high URIfreq group supports the idea that this pipeline can be used to identify children who are at a higher risk for upper respiratory infection and therefore pulmonary exacerbations that drive rapid lung function decline. Interestingly, we observed significant differences in richness of the communities associated with the “high” URIfreq group, but no difference in evenness of beta diversity. These data likely indicate a complex relationship between intestinal gut microbiota composition and specific airway clinical outcomes.

Because previous literature has shown that the microbes in the gut can alter systemic inflammatory profiles^6,10,13,14,16,18,26,28,29^, we sought to apply this machine learning predictive pipeline to markers of systemic inflammation. Again, our model was able to correlate stool microbial composition with the High NLR group (i.e., the highest systemic inflammation) with relatively low error. Interestingly, we and others have shown that *Bacteroides* is depleted in the gut of infants and cwCF ^1,2,6,21,26^. While the levels of *Bacteroides* is low compared to healthy controls for all our samples ^7^, we observed that the High NLR group had the highest relative abundance of *Bacteroides* (**Figure 4C**). This somewhat surprising finding indicates that other microbes might be driving this clinical outcome, a finding consistent with the Impact Factor analysis shown in **Figure 4B**. Additionally, previous studies have shown a positive relationship between *Bacteroides eggerthii* and NLR in adults ^24^, indicating that if this species of *Bacteroides* is present, it might be driving the observed outcome. Furthermore, the High NLR group also shows an increase of 200% in *Clostridium* relative abundance, a genus known to be elevated in CF mice ^28^. Together, these data could be used to generate hypotheses to explore the mechanisms whereby these microbial communities may be driving the observed clinical outcomes.

One priority of CF research is to decrease the high burden of medical procedures that cwCF experience. Here, we show proof of concept that stool samples, collected in diapers by parents or physicians in regular clinical visits, could identify children with the highest likelihood of negative respiratory outcomes. Analyzing stool samples could replace the use of uncomfortable oropharyngeal sampling or respiratory lavage samples for diagnostics. By training a predictive model on stool microbiota composition, we aim to predict likelihood of cwCF having frequent upper respiratory infections and/or showing high levels of systemic inflammation. Applying such approaches to CF would build on previous use of such tools in the context of other diseases ^23–25,30^. For example, machine learning applied to predictive modeling was able to prevent 30% of inappropriate antibiotic therapies in a randomized control trial ^24^. Thus, the feasibility of using stool samples to predict clinical outcomes has promise in CF.

## Materials and Methods

### Microbial Sequencing of Samples

As described previously ^11^, stool in diapers was collected by parents of cwCF and stored in home freezers; upon arrival in the lab, these diaper samples were stored frozen at −80°C until processing. As reported previously, all DNA was extracted from 100 mg of thawed stool sample using the Zymo Quick-DNA Fecal Microbe miniprep kit (Cat. No. D6010) as per the manufacturer’s instructions. Illumina targeted sequencing of the V4-V5 region of the 16S rRNA gene was performed at the Marine Biological Laboratory in Woods Hole, MA. Adapter sequences were removed using cutadapt (v 2.4). All samples were further processed through the DADA2(v 1.22.0) pipeline for quality filtering and trimming (trimmed at 220 and 220 for forward and reverse reads, respectively) and merging of paired end reads to produce an amplicon sequence variant (ASV) table. ASVs were classified with taxa from the SILVA database (vSILVA_SSU_r138_2019) using the DECIPHER (v 2.22.0) pipeline. All sequence reads can be found in GenBank with CF cohort sequences found under accession number PRJNA170783 (SRP014429).

### Analysis of Microbial Communities

All code is available on GitHub (https://github.com/GeiselBiofilm). Relative abundance, alpha diversity, and beta diversity measured were calculated and plotted for each sample using the package phyloseq (v1.38.0) and ggplot2 (v3.3.5). Each sample’s ASV counts were normalized to percentage out of the sum of counts for that sample to obtain a relative abundance. The relative abundances of the top 10 genera for each classification (age, URIfreq, or NLR group) were calculated by averaging the relative abundance of all samples that belonged within that group. Phyloseq was used to plot alpha diversity metrics (Chao-1 and Shannon), and beta diversity calculated by Bray-Curtis distances. Analysis and permutation multivariate analysis of variance (ANOVA, PERMANOVA) with multiple comparisons were applied to determine statistical significance for alpha and beta diversity, respectively.

### Predictive modeling of clinical data using bacterial populations in stool samples

Three random forests classification models were trained with the relative abundance of ASVs (at the genus level) from stool samples from cwCF. The R package randomForest (v 4.7.1) was used with ntree = 10000, using default mtry of sqrt(p) where p is the number of input variables (or microbial genera) for building the tree. For the age prediction model, samples were labeled with the age of the cwCF at which the sample was collected. Each sample was therefore classified as belonging to 1 of 5 groups: <1, 1, 2, 3, or 4 (Age (years)). 21 samples were randomly selected from each age group in order to train the age prediction model. For the URIfreq model, we first calculated the total URI per patient and normalized by age to determine URIfreq per patient. Each sample was then classified as coming from a patient with Low, Medium, or High URIfreq. For the NLR model, we calculated a neutrophil to lymphocyte ratio by dividing absolute neutrophil counts by absolute lymphocyte counts. These values were then merged with stool samples from cwCF that were collected within the same 3-month period. Each sample was then classified as having a Low, Medium, or High NLR associated with the sample. We evaluated model performance based on the OOB per model. We identified taxa with the highest informative impact by reviewing the random forest Mean Decrease Accuracy and Mean Decrease Gini values. The top 10 microbial taxa are listed in the supplemental figures, and subset to the top 5 microbial taxa in the main figures for simpler visualization and discussion.

## Acknowledgements

This work was supported by grants OTOOLE19GO and 003037G221 from the Cystic Fibrosis Foundation to GAO, MADAN596389 from the CFF to JCM, T32 HL134598-01 from the NIH and the Innovation PhD Program at Dartmouth to RAV, and T32-AI007363. Additional support was provided by DartCF (NIH grant P30-DK117469) and the Cystic Fibrosis Foundation Research Development Program (STANTO19R0).

