## Supplemental Figures for "Predicting Clinical Outcomes in Infants With Cystic Fibrosis From Stool Microbiota using Random Forest Algorithms"

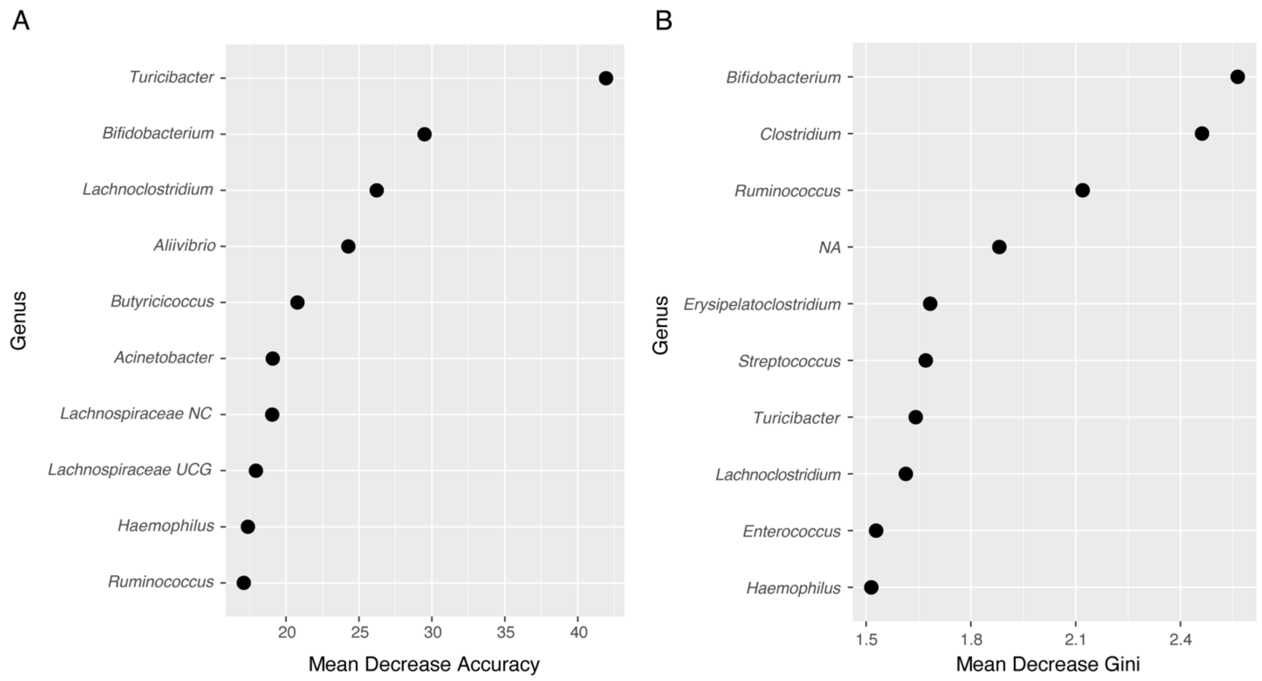


**Supplemental Figure 1. Age Model Impact Factors and Node Purity.** (A) Mean Decrease Accuracy summary plot for the top 10 stool microbial taxa important for classifying samples in the age model. B) Mean Decrease Gini summary plot for the top 10 stool microbiota with highest node purity.

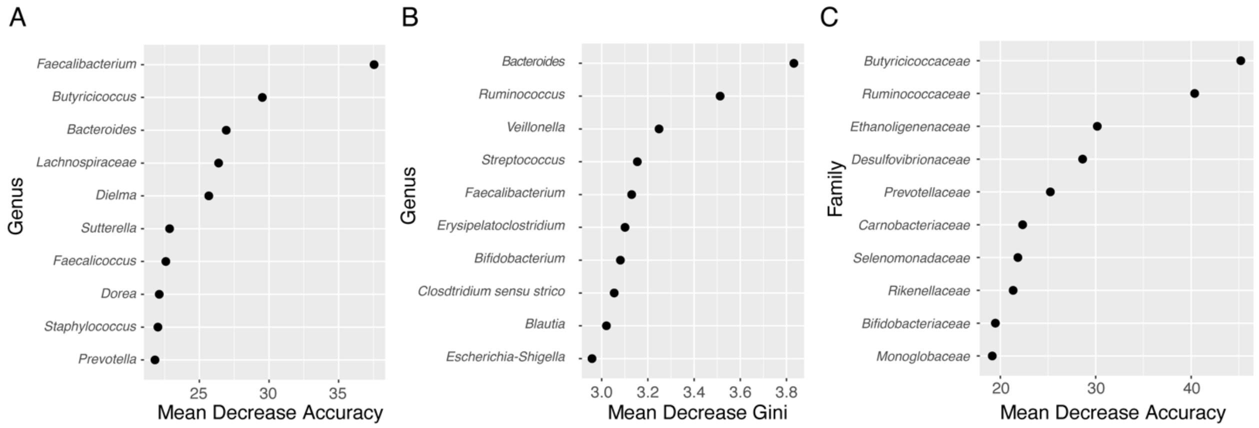


**Supplemental Figure 2. URIfreq Model Impact Factors and Node Purity.** (A) Mean Decrease Accuracy summary plot for the top 10 stool microbial taxa important for classifying samples in the URIfreq model. B) Mean Decrease Gini summary plot for the top 10 stool microbial taxa with highest node purity in the URIfreq model. (C) Mean Decrease Accuracy summary plot for the top 10 stool microbial taxa important for classifying samples in the URIfreq model trained on ASVs combined at a phylum level.

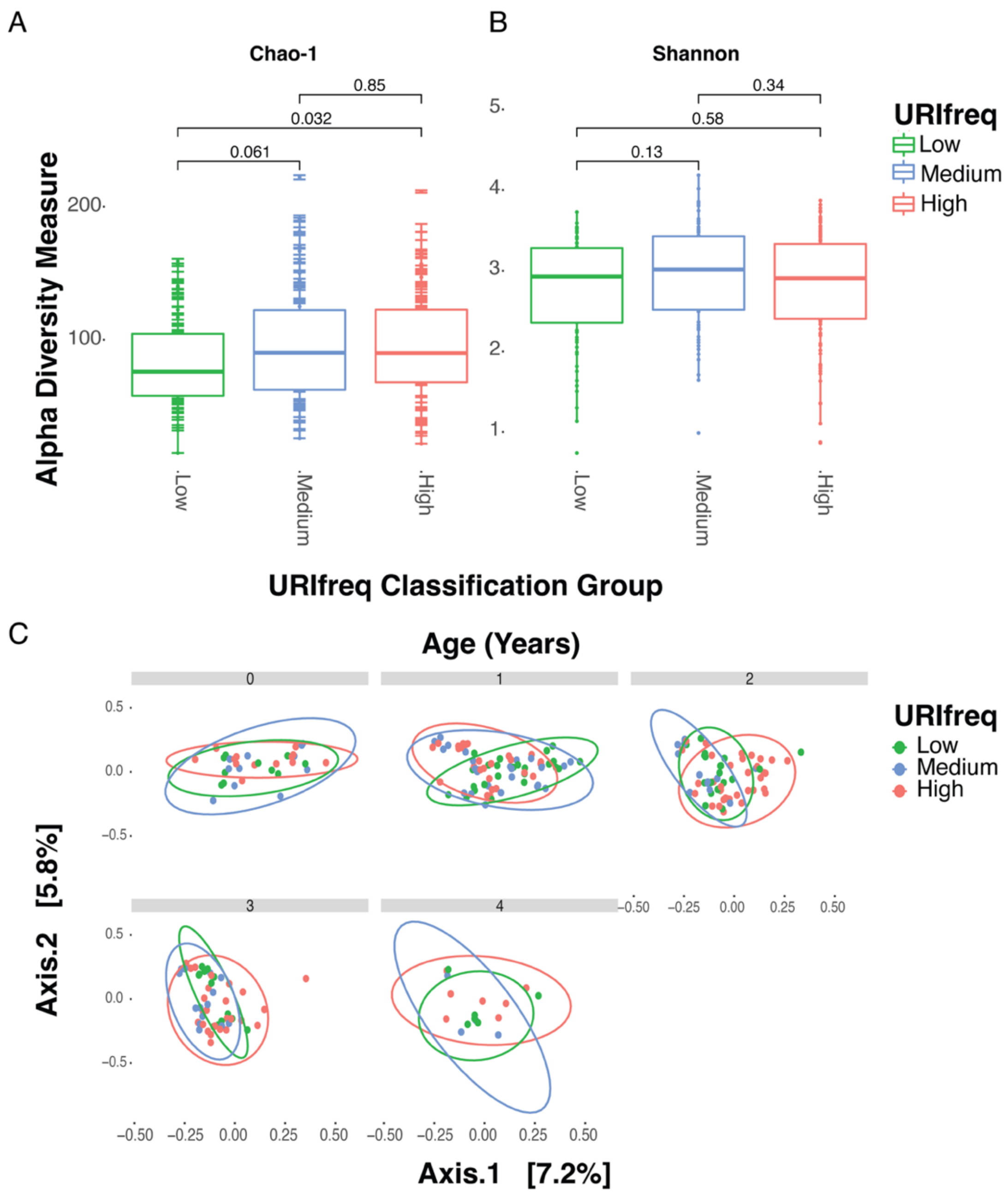


**Supplemental Figure 3. Association of microbial composition and URI**. (A) Chao-1 alpha diversity of stool microbiota in URIfreq groups. There is a significant increase in Chao-1 alpha diversity between the Low and High groups of URIfreq. (B) Shannon-alpha diversity of stool microbiota in URIfreq groups. Statistically determined by Wilcoxon rank sum with FDR correction for multiple testing. (C) PCoA plot depicting beta diversity distances between URIfreq groups at each year of life. Composition of URIfreq groups is systematically different when the interaction of age is accounted for (P==0.009), as determined by PERMANOVA. However, factors other than URIfreq and age seem to drive most of the differences in composition because these factors explain little of the variability in composition (R2 = 0.011).

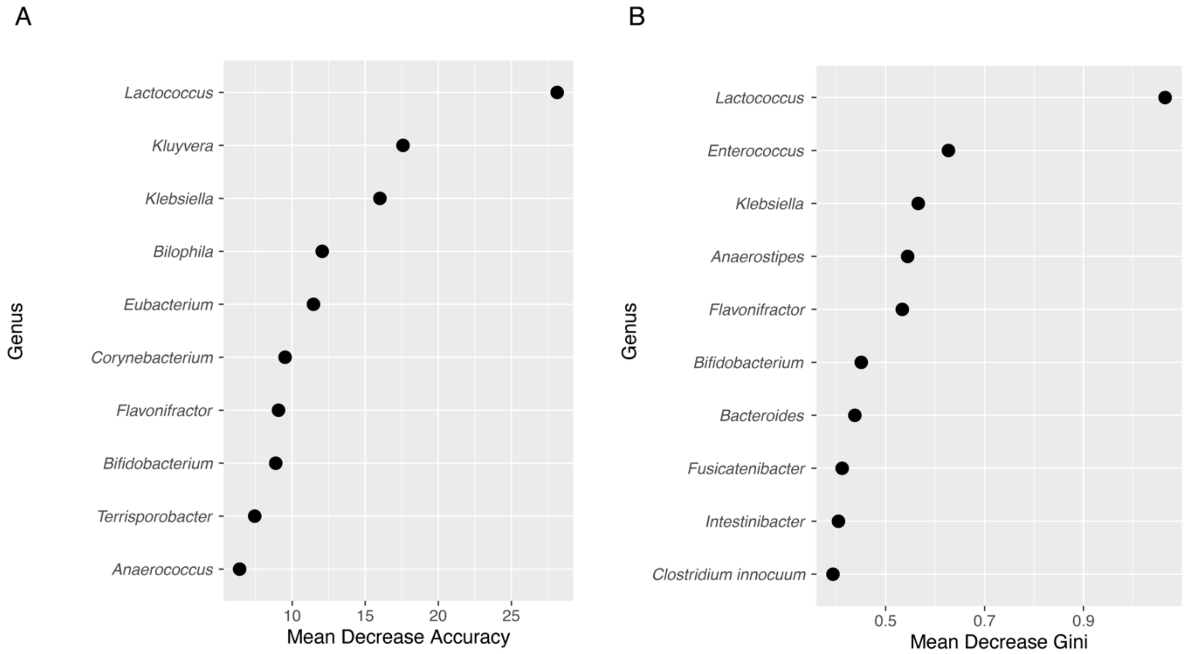


**Supplemental Figure 4. NLR Model Impact Factors and Node Purity.** (A) Mean Decrease Accuracy summary plot for the top 10 stool microbial taxa important for classifying samples in the NLR model. B) Mean Decrease Gini summary plot of the top 10 microbial taxa with highest node purity in the NLR model.

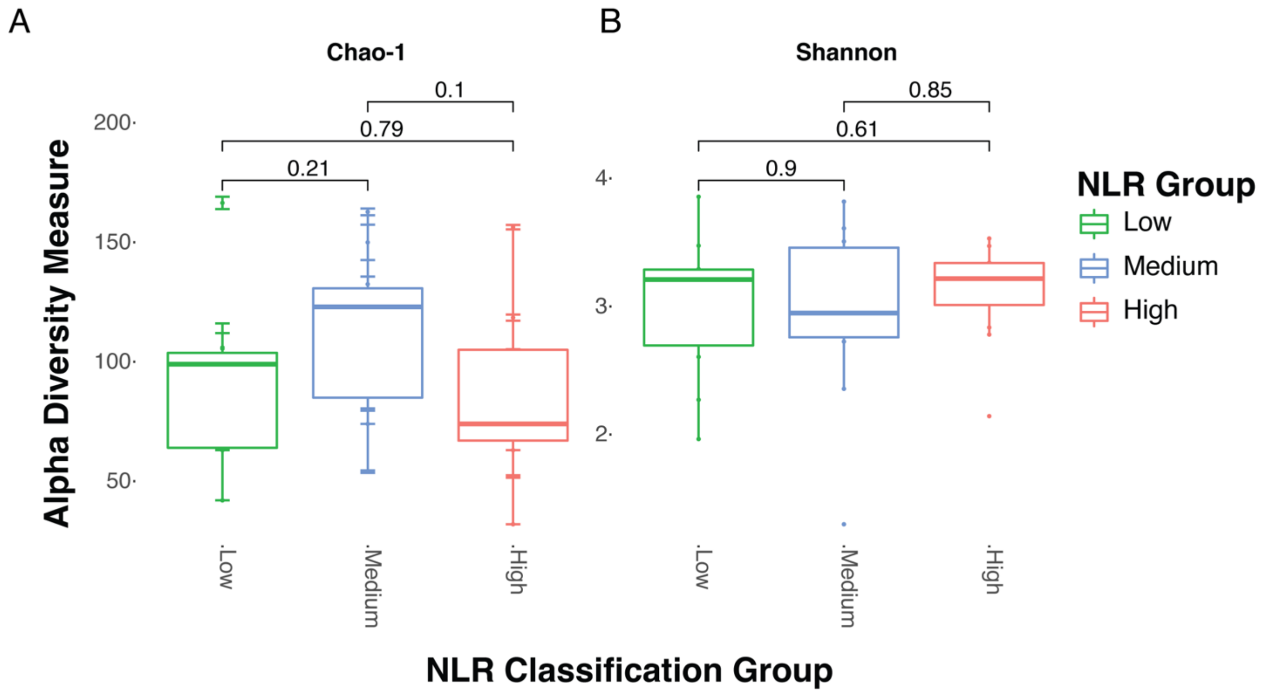


**Supplemental Figure 5. Association of microbial composition and NLR**. (A) Chao-1 alpha diversity of stool microbiota in NLR groups. (B) Shannon-alpha diversity of stool microbiota in NLR groups. Statistically determined by Wilcoxon rank sum with FDR correction for multiple testing.
